# The weakly electric fish, *Apteronotus albifrons*, avoids hypoxia before it reaches critical levels

**DOI:** 10.1101/2020.05.14.095398

**Authors:** Stefan Mucha, Lauren J. Chapman, Rüdiger Krahe

## Abstract

Anthropogenic environmental degradation has led to an increase in the frequency and prevalence of aquatic hypoxia (low dissolved-oxygen concentration, DO), which may affect habitat quality for water-breathing fishes. The weakly electric black ghost knifefish, *Apteronotus albifrons*, is typically found in well-oxygenated freshwater habitats in South America. Using a shuttle-box design, we exposed juvenile *A. albifrons* to a stepwise decline in DO from normoxia (>95% air saturation) to extreme hypoxia (10% air saturation) in one compartment and chronic normoxia in the other. Below 22% air saturation, *A. albifrons* actively avoided the hypoxic compartment. Hypoxia avoidance was correlated with upregulated swimming activity. Following avoidance, fish regularly ventured back briefly into deep hypoxia. Hypoxia did not affect the frequency of their electric organ discharges. Our results show that *A. albifrons* is able to sense hypoxia at non-lethal levels and uses active avoidance to mitigate its adverse effects.

**Summary:** The weakly electric knifefish, *Apteronotus albifrons*, avoids hypoxia below 22% air saturation. Avoidance correlates with increased swimming activity, but not with a change in electric organ discharge frequency.

## 1. Introduction

All water-breathing fishes depend on dissolved oxygen (DO) for their long-term survival (Kramer, 1984). In many aquatic ecosystems, DO concentration fluctuates naturally and can reach critically low levels due to water stratification and high temperatures as well as biological decomposition and respiration processes (Diaz, 2001; Graham, 1990; Kramer, 1984), a condition called aquatic hypoxia. Natural hypoxia is particularly widespread in tropical freshwaters where high water temperature elevates organic decomposition and reduces oxygen solubility. In addition, recent anthropogenic influences such as global climate change and eutrophication of water bodies have led to an increase of frequency and severity of hypoxic events in oceanic and coastal regions (Breitburg et al., 2018; 2018; Diaz and Rosenberg, 2008; Goldberg, 1995; Pörtner, 2001; Pörtner and Knust, 2007; Pörtner and Peck, 2010; Schmidtko et al., 2017) as well as freshwater lacustrine systems (Jenny et al., 2016a; Jenny et al., 2016b). Many fishes respond to hypoxia by migrating to better oxygenated habitats if those are available (Bell and Eggleston, 2005; Brown et al., 2015; Crampton, 1998; Pihl et al., 1991). Such avoidance behaviour can provide individuals with the flexibility to mitigate hypoxic stress without the immediate need for physiological or biochemical adjustments, though this is largely speculative. Furthermore, not all fish species show active avoidance behaviour (Cook et al., 2011), and some hypoxia-tolerant species even actively seek hypoxic zones as refuges from predators (Anjos et al., 2008; Chapman et al., 2002; Vejřík et al., 2016). To broaden our understanding of hypoxia avoidance behaviour in fish we subjected a Neotropical weakly electric fish to oxygen choice experiments. We selected a species that is reported as sensitive to hypoxia and therefore likely to exhibit active avoidance behaviour. To our knowledge, this is the first study of a weakly electric fish in a behavioural hypoxia avoidance experiment.

The black ghost knifefish, *Apteronotus albifrons*, belongs to the gymnotiform weakly electric fishes, a group that constitutes a major food web component in many floodplains of the Amazon and Orinoco basins (Crampton, 1996; Lundberg et al., 1987). Weakly electric fish generate an electric field around their body by discharging a specialized electric organ. *Apteronotus albifrons* produces wave-type electric organ discharges (EODs): continuous, quasi-sinusoidal EODs with frequencies between 800 and 1200 Hz (Crampton and Albert, 2006; Hopkins, 1976). By sensing perturbations of the electric field, weakly electric fish are able to navigate and locate objects in dark and turbid waters (Lissmann and Machin, 1958) and communicate with conspecifics (Heiligenberg, 1989). Their EODs are easy to measure, which makes them particularly well suited to research on the energetics of sensation and communication (e.g. Julian et al., 2003; Markham et al., 2016; Moulton et al., 2020; Reardon et al., 2011; Salazar et al., 2013). As apteronotid fish naturally occur in well oxygenated habitats (Crampton, 1998), it has been suggested, that they are not able to tolerate hypoxia. The aim of our study was to find out how hypoxia affects the swimming behaviour and the active electric sense of *A. albifrons*. We quantified swimming behaviour and EOD frequency while exposing fish to progressive hypoxia in a shuttle-box choice chamber and offering a normoxic refuge at all times. We hypothesized that *A. albifrons* will begin to avoid hypoxia at moderate DO levels as part of their natural respiratory strategy. Based on a study of the closely related brown ghost knifefish, *Apteronotus leptorhynchus*, which only found a small decrease in EOD frequency under hypoxic stress (Reardon et al., 2011), we hypothesized that *A. albifrons* will not modulate their EOD frequency while experiencing hypoxia.

## 2. Materials and Methods

### Experimental animals and housing conditions

We used farm-bred *Apteronotus albifrons* (Linnaeus, 1766) obtained from a commercial supplier (AQUAlity Tropical Fish Wholesale, Inc., Mississauga, Ontario, Canada). Experiments were performed with 16 individuals with a mean body mass of 3 g (range: 1.7 – 4.2 g), a mean standard body length (SBL) of 8.9 cm (range: 7.6 – 10.4 cm), and an electric organ discharge (EOD) frequency of 807 – 1151 Hz at 26°C. Sexually mature *A. albifrons* typically have a SBL of 14 – 30 cm and a body mass of at least 20 g (Dunlap and Larkins-Ford, 2003; Nelson and MacIver, 1999; Serrano-Fernández, 2003). Thus, it is likely that most, if not all, of our experimental animals were sexually immature, and we did not distinguish fish by sex for data analysis. This was confirmed by gonadal inspection in one case. Fish were housed in tanks of 75 L in groups of 3 – 4 individuals per tank. Individual fish were separated with plastic mesh tank dividers, and each fish had access to one PVC tube as shelter. The water temperature averaged 25.7°C (range: 25.4 - 25.9°C), conductivity 200 μS (190 - 210 μS), and pH 7.1 (6.8 – 7.3). Normoxic air saturation levels (>95%) were maintained by bubbling air into the tanks. Fish were kept at a 12:12 h light:dark photoperiod and were fed daily a small amount of frozen bloodworms (chironomid larvae, Hikari Sales USA, Inc., Hayward, California, USA). Controlled conditions were maintained for a minimum of two weeks before the start of experiments. All procedures were approved by the McGill University Animal Care Committee (protocol # 5408).

### Hypoxia avoidance setup

We used a shuttle-box dissolved oxygen choice chamber (Loligo Systems Inc., Denmark) to quantify hypoxia avoidance behaviour (Fig. 1). The choice chamber consisted of two circular compartments (each 50 cm in diameter) connected by a central passage (W = 8.5 cm, L = 14 cm). A PVC tube (L = 15 cm, inner diameter 2.6 cm) was placed symmetrically in both compartments as shelter to minimize stress and to reduce arbitrary swimming activity. The two compartments received water from separate buffer tanks where the dissolved oxygen (DO) was controlled by bubbling air or nitrogen gas into the water. DO was measured before the water entered the compartments with a galvanic oxygen probe (MINI-DO, Loligo Systems). Water exchange between choice chamber and buffer tanks was maintained with aquarium pumps (Universal Pumpe 1048, EHEIM GmbH & Co.KG, Germany). Water temperature was maintained via silicon rubber heating mats (OMEGA Engineering, Inc., USA) that were wrapped around the buffer tanks and controlled by a thermostat (Inkbird Tech C.L., China) with a submerged temperature sensor placed in the passage between the compartments. Circular acrylic glass lids were submerged in the choice chamber ca. 1 cm below the water surface to reduce the diffusion of atmospheric oxygen into the water and to prevent fish from accessing the surface during trials.

**Fig. 1:**
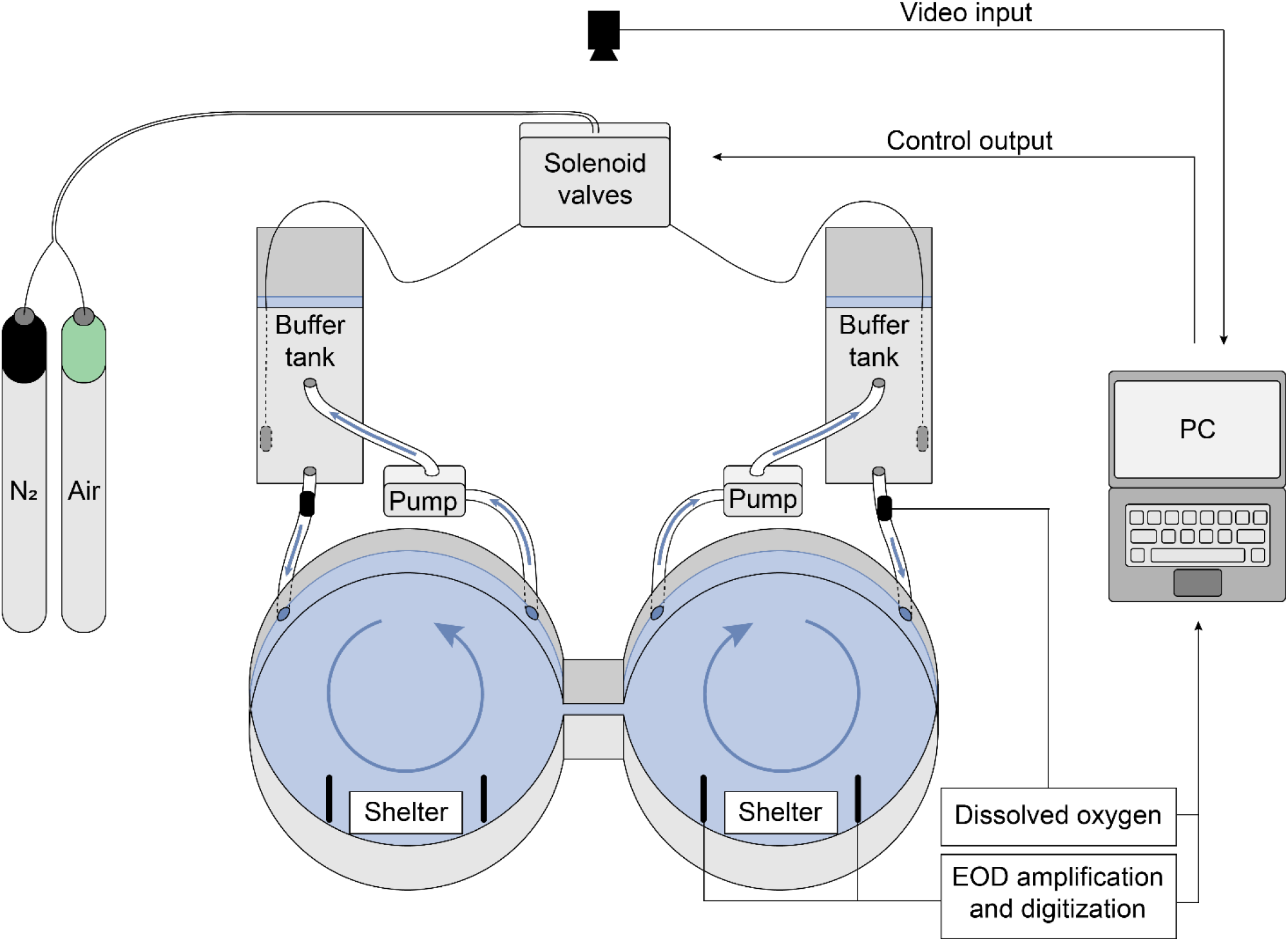
Schematic of shuttle-box oxygen choice chamber. Blue arrows indicate water flow in the compartments and between choice chamber and buffer tanks. Plexiglas lids, heating system and grounding electrode are not shown.

Fish position was recorded with a camera (UI-1640-SE-C-GL, IDS Imaging Development Systems GmbH, Germany) mounted above the shuttle-box. ShuttleSoft software (ver. 2.6.4, Loligo Systems) was used to log fish position and DO and to control air saturation in the buffer tanks via a DAQ-M device (Loligo Systems Inc., Viborg, Denmark), which operated solenoid valves at the gas tubing. EODs were measured via submerged carbon rod electrodes in the choice chamber. Two electrodes were placed in each compartment near the PVC tube that served as a shelter, and one grounding electrode was placed in the passage between the compartments. The choice chamber was set up in an isolated room to minimize disturbance.

### Hypoxia avoidance trials

Trials were conducted at water parameters resembling housing conditions (conductivity of 200 μS and pH of 6.9 – 7.3) with a total water volume of 60 L. Due to varying room temperatures, water temperature at the initiation of the trials varied between 25.4 and 25.9°C. During trials, temperature decreased on average by 0.15°C (0-0.4°C) due to room ventilation.

Fish were fasted for 36 h prior to experiments to ensure a post-absorptive state. For each trial, one fish was introduced into the choice chamber in the afternoon and left for 16 h to acclimate overnight. The side of introduction was chosen randomly, and the fish could freely shuttle between both compartments throughout the experiment. Water was aerated until the start of trials, and measurement devices were calibrated to 100% air saturation before each trial. Trials started in the morning at 9:30 h (30 min after the onset of the light photoperiod). Both compartments were maintained at >95% air saturation for 40 min to record baseline behaviour at normoxia. Each of the 16 fish exhibited a pronounced preference for one of the two compartments during baseline controls. We subsequently induced stepwise hypoxia in the compartment of the choice chamber where the fish preferred to stay while maintaining water in the non-preferred compartment at high DO levels (> 80% air saturation). DO concentration was incrementally lowered to the following air saturation levels: 70%, 50%, 30%, 25%, 20%, 15%, and 10%. Each DO concentration was maintained for 10 min followed by a 10 min decrease to the next lower concentration (Fig. 2A). After the lowest DO concentration was reached, the hypoxic compartment was reoxygenated, and data acquisition was continued for 20 min. The total trial duration (baseline + hypoxia induction + reoxygenation) was 200 min. Upon completion of a trial, the fish was weighed and its SBL measured.

**Fig. 2:**
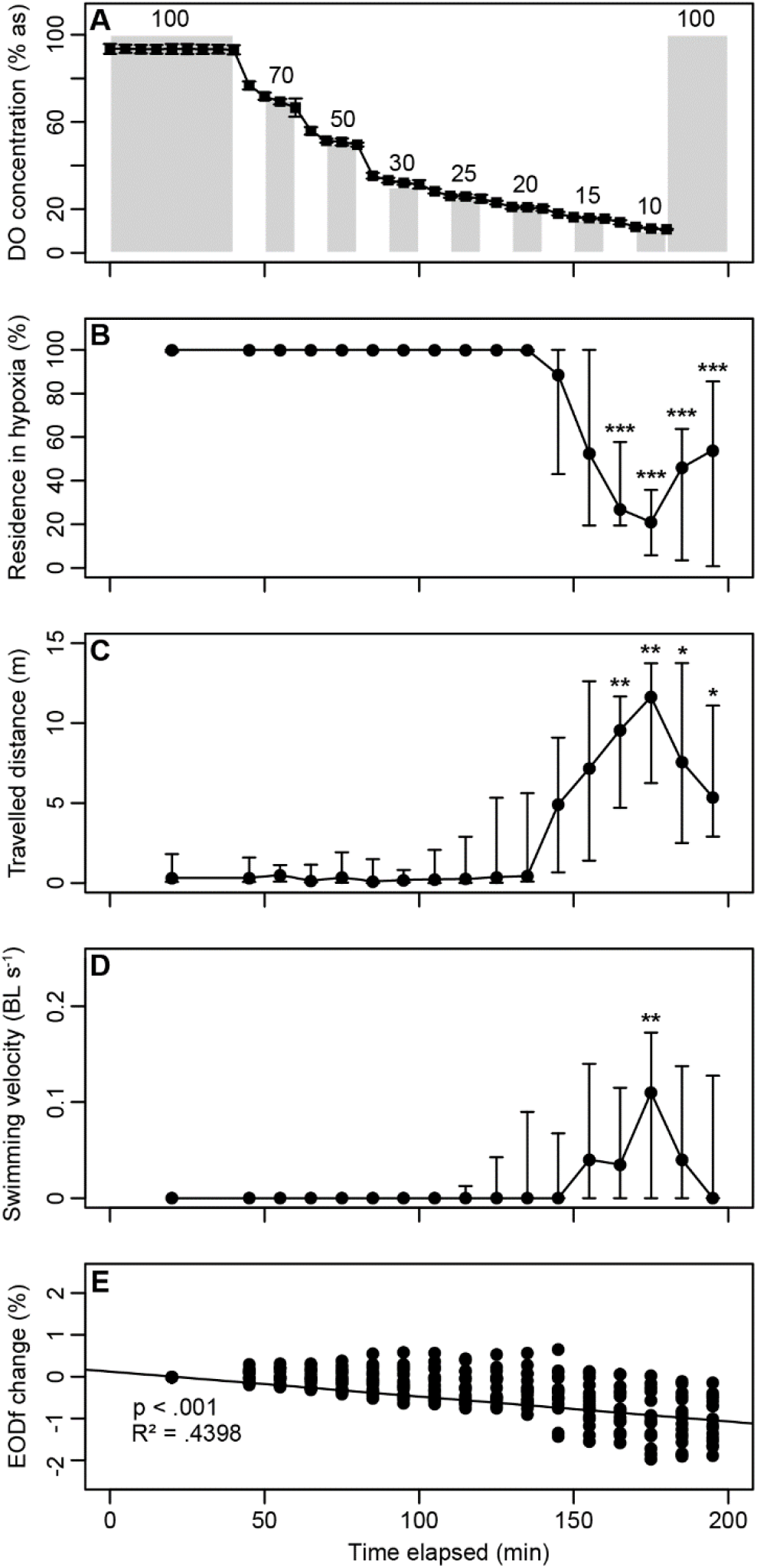
Behavioural responses of *A. albifrons* during hypoxia avoidance trials. (A) DO concentrations during a control trial with no fish. Grey bars represent the target DO concentration, black circles represent control measurements. (B) Percentage of time spent in the hypoxic compartment. We induced hypoxia in the compartment where the fish preferred to stay. (C) Distance travelled in the entire choice chamber. (D) Swimming velocity in the entire choice chamber. (E) Linear regression of EOD frequency change as percentage change from baseline EOD frequency based on LMM. Circles represent median values, vertical bars represent first and third quartile, values from the first 40 min were pooled as normoxic baseline behaviour, asterisks indicate statistically significant differences from normoxic baseline behaviour (pairwise Wilcoxon rank-sum tests with Holm-Bonferroni correction of p-values, p < 0.05 *, p < 0.01 **, p < 0.001 ***).

### Data acquisition and processing

During each trial, the fish position was tracked from above based on image contrast. X and Y coordinates, distance moved (cm), swimming velocity (cm s^−1^), and air saturation (%) were logged every second. The log file was processed with Microsoft Excel^®^2010 (Microsoft Corp., Redmond, Washington, USA) and R (ver. 3.2.5, https://www.r-project.org). Electrical EOD recordings were band-pass filtered (300 Hz – 5 kHz) and amplified (1000x gain, A-M Systems Model 1700, USA). Signals were then digitized with a sample rate of 20 kHz (National Instruments USB-6211, USA) and saved on a computer using custom-written Matlab programs (The MathWorks, Inc., USA).

### Statistical analyses

All statistical analyses were performed with R (ver. 3.2.5, https://www.r-project.org). Raw data from video tracking and processed datasets used for statistical analyses are available online (uploaded to figshare repository, link will be made available upon acceptance).

### Side preference during normoxic baseline recordings

Side preference was tested with a two-sided single-sample Wilcoxon rank-sum test on residence time in the preferred compartment against the null hypothesis that fish would spend 50% of the time in each of the compartments (= no preference). As fish did not tend to rest in the passage between compartments, we ignored this possibility for this test.

### Swimming behaviour

Residence time in hypoxia (% of time spent in the hypoxic compartment), average swimming speed (body lengths per second, BL s^−1^), and distance moved (m) were summarized as medians over the 40 min normoxic baseline period and each following 10 min interval of the trial.

Residence time in the hypoxic compartment, swimming speed and distance were tested for significant changes throughout the trial using Friedman’s rank-sum test with experimental time as independent variable. In case of a significant result, this was followed by pairwise Wilcoxon rank-sum tests with Holm-Bonferroni correction of p-values to identify the experimental time at which a significant deviation from normoxic baseline recordings occurred.

Due to water exchange between the choice chamber and buffer tanks, there was a constant circular water current in the compartments. Swimming speed and distance were not corrected for water current; rather, these metrics are used to indicate changes of swimming activity, such as stationary behaviour vs. exploration/avoidance.

### Electric organ discharges

EOD frequency was extracted from recordings using custom written routines in Matlab R2017a (The MathWorks, Inc., Natick, Massachusetts). Recorded signals were Fourier-transformed, and the frequency with the highest power spectral density estimate (frequency resolution 0.076 Hz) was picked as the EOD frequency for every second of the recording. Median EOD frequency over the 40 min baseline period and each following 10 min interval of the trial was calculated for each fish. To account for individual differences in the baseline EOD frequency of each fish, values were normalised as percent change from normoxic baseline values for each of the 10 min intervals following baseline recordings. To test for an effect of hypoxia on EOD frequency, we used a random-slope linear mixed-effect model (LMM) with change of median EOD frequency as dependent variable. Based on AIC score, the best fit was achieved by including the interaction of inversed DO concentration with residence time in hypoxia (i.e. the lower the DO concentration in which the fish stayed, the higher the interaction term) and experimental time as fixed effect and fish ID as random effect. The intercept of the LMM was set to zero.

EOD amplitude was strongly affected by the position and orientation of the fish relative to the recording electrodes. As we could not always determine the exact fish position and orientation (e.g. when fish were in their shelters or swimming in the passage between compartments), we excluded EOD amplitude from our analysis.

### Hypoxia avoidance threshold

We quantified the threshold for hypoxia avoidance by modelling the correlation between residence time in hypoxia and DO concentration with a modified version of the program for P_crit_-determination by Yeager and Ultsch (Yeager and Ultsch, 1989). The program estimates the best fit of two linear regressions to a dataset, iteratively minimizing their residual sum of squares. Two LMMs with random intercepts were calculated with residence time in the hypoxic compartment as the dependent variable. DO concentration was included as fixed effect, and fish ID was included as a random effect. The hypoxia avoidance threshold was defined as the DO concentration at which the regression lines of both LMMs intersected. Conditional and marginal R^2^ values of both LMMs were calculated based on the method by Nakagawa and colleagues (Nakagawa et al., 2013). T-test statistics and p-values for the null hypothesis of zero correlation between residence time in hypoxia and DO were calculated with degrees of freedom obtained through Satterthwaite approximation. The R code for these procedures was adapted from the rMR package (Moulton, 2018).

### Repeatability trials

To test the repeatability of our experimental protocol and results, hypoxia avoidance trials were repeated after 4 weeks with five fish. Trials were conducted as described above, and residence time in the hypoxic compartment was tested for differences between the first trial and the repeatability trial using a two-way repeated measures ANOVA with DO concentration as between-subject effect, and experimental day as within-subject effect.

## 3. Results and Discussion

### Apteronotus albifrons show pronounced side preference and stationary behaviour at normoxia

During normoxic baseline recordings, all 16 individuals showed a pronounced preference for one compartment of the shuttle-box choice chamber over the other (Wilcoxon single sample rank-sum test, p < 0.001) with 10 fish spending the whole baseline period exclusively on one side and no fish spending less than 79% of the time on one side. Among all 16 fish, the two compartments were chosen 8 times each, indicating that there was no bias to either side of the choice chamber. During this period, fish predominantly rested in the PVC tubes that were provided as shelters. We subsequently induced stepwise hypoxia in the compartment of the choice chamber where the fish preferred to stay.

### Increased locomotor activity drives hypoxia avoidance at safe oxygen levels

Moderate hypoxia above 20% air saturation did not significantly affect swimming behaviour. Fish predominantly rested in their shelters and showed only small deviations from normoxic baseline behaviour. Below 20% air saturation, swimming activity increased and fish spent less time in the hypoxic compartment with significant deviations from baseline recordings below 15% air saturation (Wilcoxon rank-sum test with Holm-Bonferroni *post-hoc* correction, p < 0.05, Fig. 2B-D, Table S1, S2). To determine the threshold for the onset of hypoxia avoidance, we modelled the impact of hypoxia on residence time in the hypoxic compartment using two linear mixed-effect models (LMMs) with random intercepts and slopes (Fig. 3, Table S3). Based on this method, we identified the threshold for the onset of hypoxia avoidance behaviour at the intersection of both linear regressions at 22% air saturation. Above the threshold, air saturation had no significant effect on the residence time in the hypoxic compartment (adjusted marginal R^2^ = 0.02, p = 0.075). Below the threshold, air saturation significantly affected residence in the hypoxic compartment (adjusted marginal R^2^ = 0.307, p < 0.001). In repeatability trials, residence times in the hypoxic compartment did not differ significantly from original trials (ANOVA: F = 0.114, p = 0.753, Table S4).

**Fig. 3:**
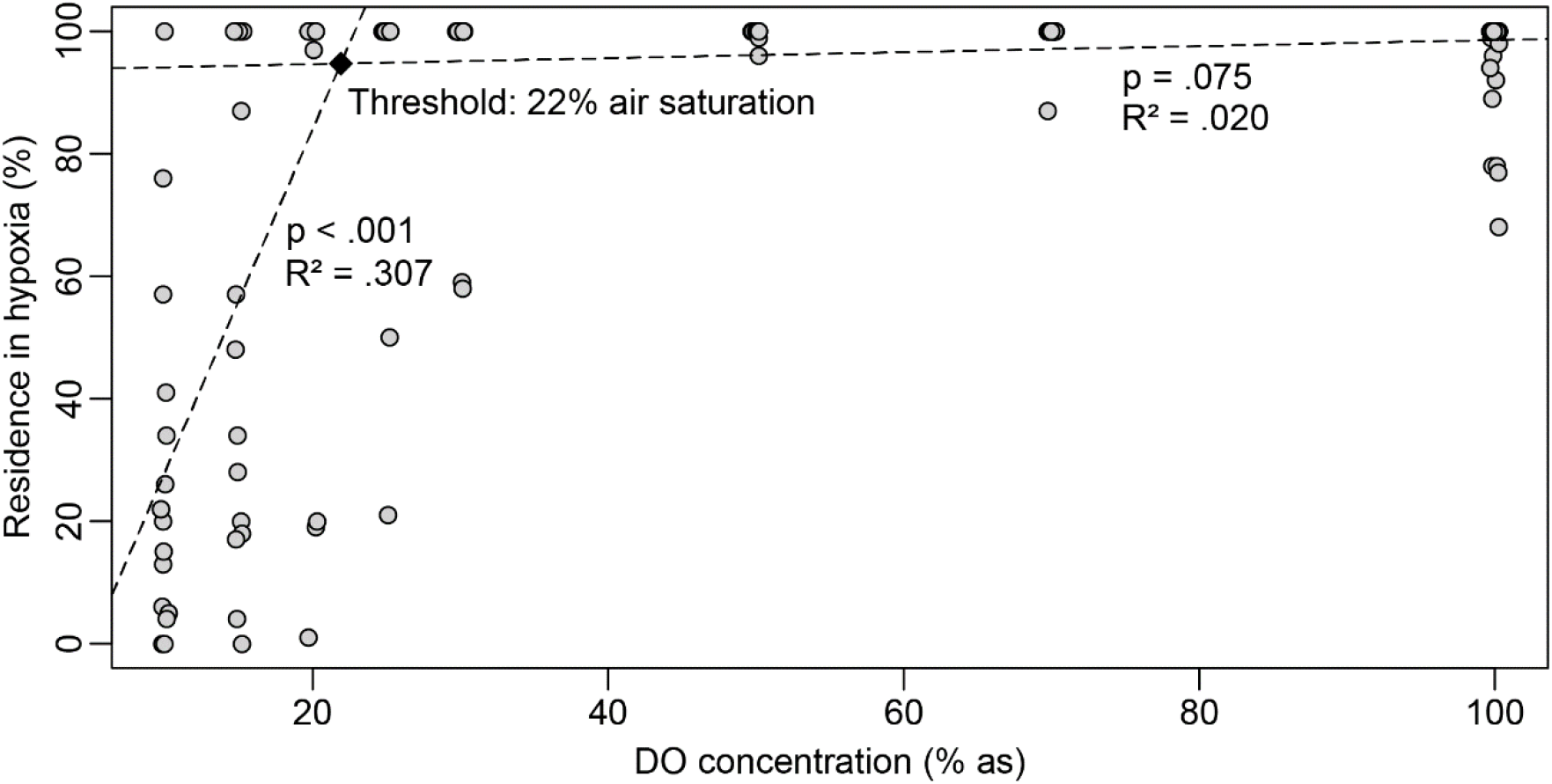
Residence time in the hypoxic compartment as function of the dissolved oxygen in % air saturation. Grey circles represent the percentage of time that individual fish spent in the hypoxic compartment at each air saturation that was established in this compartment (n = 16 fish, points are jittered along the x-axis to reveal overlapping measurements), dashed lines represent linear regressions based on LMMs, the black diamond at their intersection indicates the computed hypoxia avoidance threshold, R^2^ = adjusted marginal R^2^, p = probability of zero correlation between air saturation and residence in hypoxia.

These results show that *A. albifrons* use an active hypoxia avoidance strategy that follows a threshold dynamic with little or no effect of moderate hypoxia and a strong effect of deep hypoxia on swimming behaviour. The avoidance threshold lies above the threshold for aquatic surface respiration for *A. albifrons* (ASR_50_, the oxygen level at which fish spend 50% of their time engaged in breathing water from the surface film), estimated as 18.3% (Vassileva, Krahe and Chapman, unpublished data). The threshold for hypoxia avoidance in *A. albifrons* also falls well above the critical oxygen tension (P_crit_) for the closely related species *Apteronotus leptorhynchus* and the distantly related gymnotiform *Eigenmannia virescens*. Pcrit, the oxygen partial pressure below which the oxygen consumption of the fish switches from oxygen regulation to oxygen conformation, was estimated as 10.5% air saturation for *A. leptorhynchus* and 7.1% for *E. virescens* (Reardon et al., 2011). The early onset of an avoidance reaction likely provides *A. albifrons* with the flexibility to seek better oxygenated areas before hypoxic stress impairs its physiology. This is consistent with anecdotal observations that wave-type gymnotiform fishes are typically found in habitats with a DO concentration of 40% - 60% air saturation and avoid swimming into hypoxic or anoxic waters (Crampton, 1998). The active response to hypoxia described here is similar to the behavioural responses of other species, such as red hake (*Urophycis chuss*, Bejda et al., 1987), tuna (*Katsuwonus pelamis* and *Thunnus albacares*, Bushnell and Brill, 1991), Atlantic herring (*Clupea harengus*, Domenici et al., 2000), weakfish (*Cynoscion regalis*, Brady et al., 2009), and rainbow trout (*Oncorhynchus mykiss*, Poulsen et al., 2011).

After leaving hypoxia, fish remained active and occasionally ventured back to the hypoxic side. This sustained change from stationary swimming to active roaming was evident in the distance travelled, which remained significantly higher than baseline values at air saturations below 15% (Wilcoxon rank-sum test with Bonferroni-Holm *post-hoc* correction, p < 0.01, Fig. 2C,D). This behaviour was somewhat unexpected as a short bout of activity would have sufficed to leave the hypoxic compartment of the choice chamber while minimizing energy expenditure and predation risk. The lasting upregulation of locomotor activity could be caused by the initial displacement from their shelter. In a natural setting, hypoxic areas are likely to be more extended than in our setup and a sustained increase of locomotor activity might be necessary to reach better oxygenated areas. Excursions into hypoxia have been found in other fish species, sometimes associated with foraging behaviour (Claireaux et al., 1995; Cook and Herbert, 2012; Herbert et al., 2011; Jones, 1952; Rahel and Nutzman, 1994; Wannamaker and Rice, 2000). Based on these findings, it has been hypothesized that hypoxia avoidance behaviour is not directly triggered by external DO concentration but rather relies on various physiological cues that imply “respiratory distress” (Cook et al., 2011; Jones, 1952). This indirect relationship between external DO concentrations and behavioural response allows for the integration of many additional cues into an avoidance response, thus increasing its flexibility in different environmental contexts. Although ultimately, the physiological need for oxygen is the driver of hypoxia avoidance, the onset of this behaviour could be dependent on the interaction of several relevant factors such as habitat cover (Hill, 1968), presence of predators (Wolf and Kramer, 1987), availability and quality of an oxygen refuge (Herbert et al., 2011), and acclimation to different oxygen regimes (Cook et al., 2013). Although *A. albifrons* are known to inhabit well-oxygenated waters, they are likely to experience hypoxia occasionally in their natural habitat. Thus the ability to venture into hypoxic waters without immediate avoidance would allow them to forage or migrate and thus could provide an important fitness benefit.

### Electric organ discharge frequency is not a part of the hypoxia mitigation strategy of A. albifrons

Median EOD frequency decreased marginally throughout hypoxia avoidance trials with the lowest values averaging to −1.02% change from normoxic baseline EOD frequency at the beginning of reoxygenation (Fig. 2E, Table S5). According to LMM estimates, this decrease resulted from a small but significant negative effect of experimental time on frequency. The most likely cause of the marginal decrease of median EOD frequency is the slight cooling of water temperature during trials, which amounted to an average decrease of 0.15°C. Assuming a Q_10_ value of 1.55 for EOD frequency (Dunlap and Ragazzi, 2015), temperature change explains a reduction of EOD frequency by 0.7%. The interaction term of residence time in hypoxia and inverted DO concentration had a negligible positive effect on frequency, indicating that fish did not reduce their EOD frequency in response to hypoxia. Although in theory, a reduction of discharge frequency might reduce the energetic cost of the electric sense and thus could be a useful means to survive hypoxia (Salazar et al., 2013), the lack of evidence that wave-type gymnotiforms employ frequency reduction as a measure to save energy suggests that wave-type gymnotiforms are unable to effectively reduce their EOD frequency, even under inescapable hypoxic stress (Crampton, 1998; Markham et al., 2009; Reardon et al., 2011). Whereas we can make no inferences about the capacity of *A. albifrons* to regulate EOD frequency under hypoxic stress, our results show that *A. albifrons* leave hypoxia before EOD frequency is affected, regardless of whether by active regulation or as a mere consequence of hypoxic stress.

Another parameter that contributes to the energetic cost of EODs is their amplitude (Markham et al., 2009; Salazar et al., 2013; Stoddard and Salazar, 2010). So far, reduction of EOD amplitude has only been found under inescapable hypoxic conditions approaching the respective P_crit_ in *A. leptorhynchus* and *E. virescens* (Reardon et al., 2011). The comparatively high hypoxia avoidance threshold of 22% air saturation suggests that *A. albifrons* avoided hypoxia well before its EOD amplitude was affected. However, as we could not reliably quantify EOD amplitude of freely swimming fish in our study, additional experiments are needed to clarify whether the DO concentrations at which fish began to leave hypoxia in our experiment have an effect on EOD amplitude.

### Conclusion and outlook

We show here that *A. albifrons* use an active hypoxia avoidance strategy that is comparable to that of other fishes with active life styles. Our results suggest that active avoidance serves to mitigate negative implications of hypoxia on sensing and physiology rather than adapting to it. These results are in line with previous studies and field observations of wave-type gymnotiforms (Crampton, 1998; Reardon et al., 2011) and suggest a low tolerance of *A. albifrons* to hypoxia below 20% air saturation. With regard to the expected increased prevalence of hypoxia in the future, this proactive avoidance strategy is likely to cause habitat shifts and a reduced abundance of *A. albifrons* in affected habitats. More hypoxia-related behavioural studies are needed for us to better understand the flexibility of behaviour in different environmental contexts and the relationship between physiological and behavioural hypoxia tolerance.

## Acknowledgements

We would like to thank Tyler L. Moulton for his assistance in implementing statistical methods and Stefan K. Hetz for guidance throughout this project.

## Author Contributions

All authors participated in the design of this study. S.M. performed all experiments and data analyses. R.K. wrote Matlab scripts for recording EODs. S.M. drafted the manuscript and all authors took part in its revision.

## Tables

See supplementary info for tables S1-S5.

## References

Bejda, A. J., Studholme, A. L. and Olla, B. L. (1987). Behavioral responses of red hake, Urophycis chuss, to decreasing concentrations of dissolved oxygen. Environ. Biol. Fish. 19, 261–268.

Bell, G. W. and Eggleston, D. B. (2005). Species-specific avoidance responses by blue crabs and fish to chronic and episodic hypoxia. Mar. Biol. 146, 761–770.

Brady, D. C., Targett, T. E. and Tuzzolino, D. M. (2009). Behavioral responses of juvenile weakfish (Cynoscion regalis) to diel-cycling hypoxia: swimming speed, angular correlation, expected displacement, and effects of hypoxia acclimation. Can. J. Fish. Aquat. Sci. 66, 415–424.

Breitburg, D., Levin, L. A., Oschlies, A., Grégoire, M., Chavez, F. P., Conley, D. J., Garçon, V., Gilbert, D., Gutiérrez, D. and Isensee, K. et al. (2018). Declining oxygen in the global ocean and coastal waters. Science. 359.

Brown, D. T., Aday, D. D. and Rice, J. A. (2015). Responses of Coastal Largemouth Bass to Episodic Hypoxia. Trans. Am. Fish. Soc. 144, 655–666.

Bushnell, P. G. and Brill, R. W. (1991). Responses of Swimming Skipjack (Katsuwonus pelamis) and Yellowfin (Thunnus albacares) Tunas to Acute Hypoxia, and a Model of Their Cardiorespiratory Function. Physiol. Zool. 64, 787–811.

Claireaux, G., Webber, D., Kerr, S. and Boutilier, R. (1995). Physiology and behaviour of free-swimming Atlantic cod (Gadus morhua) facing fluctuating salinity and oxygenation conditions. J. exp. Biol. 198, 61–69.

Cook, D. G. and Herbert, N. A. (2012). The physiological and behavioural response of juvenile kingfish (Seriola lalandi) differs between escapable and inescapable progressive hypoxia. J. Exp. Mar. Biol. Ecol. 413, 138–144.

Cook, D. G., Iftikar, F. I., Baker, D. W., Hickey, A. J. R. and Herbert, N. A. (2013). Low-O2 acclimation shifts the hypoxia avoidance behaviour of snapper (Pagrus auratus) with only subtle changes in aerobic and anaerobic function. J. exp. Biol. 216, 369–378.

Cook, D. G., Wells, R. M. G. and Herbert, N. A. (2011). Anaemia adjusts the aerobic physiology of snapper (Pagrus auratus) and modulates hypoxia avoidance behaviour during oxygen choice presentations. J. exp. Biol. 214, 2927–2934.

Crampton, W. G. R. (1996). Gymnotiform fish: An important component of Amazonian floodplain fish communities. J. Fish Biol. 48, 298–301.

Crampton, W. G. R. (1998). Effects of anoxia on the distribution, respiratory strategies and electric signal diversity of gymnotiform fishes. J. Fish Biol. 53, 307–330.

Crampton, W. G. R. and Albert, J. S. (2006). Evolution of Electric Signal Diversity in Gymnotiform Fishes: Part A. Phylogenetic Systematics, Ecology, and Biogeography. In Communication in fishes, pp. 647–731. Enfield, NH: Science Publ.

Diaz, R. J. (2001). Overview of Hypoxia around the World. J. Environ. Qual. 30, 275–281.

Diaz, R. J. and Rosenberg, R. (2008). Spreading dead zones and consequences for marine ecosystems. Science. 321, 926–929.

Domenici, P., Steffensen, J. F. and Batty, R. S. (2000). The effect of progressive hypoxia on swimming activity and schooling in Atlantic herring. J. Fish Biol. 57, 1526–1538.

Dunlap, K. D. and Larkins-Ford, J. (2003). Diversity in the structure of electrocommunication signals within a genus of electric fish, Apteronotus. J. Comp. Physiol. A Neuroethol. Sens. Neural. Behav. Physiol. 189, 153–161.

Dunlap, K. D. and Ragazzi, M. A. (2015). Thermal acclimation and thyroxine treatment modify the electric organ discharge frequency in an electric fish, Apteronotus leptorhynchus. Physiology & behavior. 151, 64–71.

Goldberg, E. D. (1995). Emerging problems in the coastal zone for the twenty-first century. Mar. Pollut. Bull. 31, 152–158.

Graham, J. B. (1990). Ecological, evolutionary and physical factors influencing aquatic animal respiration. Am. Zool. 30, 137–146.

Heiligenberg, W. (1989). Coding and processing of electrosensory information in Gymnotiform fish. J. exp. Biol. 146, 255–275.

Herbert, N. A., Skjæraasen, J. E., Nilsen, T., Salvanes, A. G. V. and Steffensen, J. F. (2011). The hypoxia avoidance behaviour of juvenile Atlantic cod (Gadus morhua L.) depends on the provision and pressure level of an O2 refuge. Mar. Biol. 158, 737–746.

Hill, L. G. (1968). Oxygen preference in the spring cavefish, Chologaster agassizi. Trans. Am. Fish. Soc. 97, 448–454.

Hopkins, C. D. (1976). Stimulus filtering and electroreception: Tuberous electroreceptors in three species of Gymnotoid fish. J. Comp. Physiol. A Neuroethol. Sens. Neural. Behav. Physiol. 111, 171–207.

Jenny, J.-P., Francus, P., Normandeau, A., Lapointe, F., Perga, M.-E., Ojala, A., Schimmelmann, A. and Zolitschka, B. (2016a). Global spread of hypoxia in freshwater ecosystems during the last three centuries is caused by rising local human pressure. Glob. Chang. Biol. 22, 1481–1489.

Jenny, J.-P., Normandeau, A., Francus, P., Taranu, Z. E., Gregory-Eaves, I., Lapointe, F., Jautzy, J., Ojala, A. E. K., Dorioz, J.-M. and Schimmelmann, A. et al. (2016b). Urban point sources of nutrients were the leading cause for the historical spread of hypoxia across European lakes. PNAS. 113, 12655–12660.

Jones, J. R. E. (1952). The Reactions of Fish to Water of Low Oxygen Concentration. J. exp. Biol. 29, 403–415.

Julian, D., Crampton, W. G. R., Wohlgemuth, S. E. and Albert, J. S. (2003). Oxygen consumption in weakly electric Neotropical fishes. Oecologia. 137, 502–511.

Kramer, D. L. (1984). The evolutionary ecology of respiratory mode in fishes: an analysis based on the costs of breathing. In Evolutionary ecology of neotropical freshwater fishes (ed. E. K. Balon and T. M. Zaret), pp. 67–80. Dordrecht: Springer Netherlands.

Lissmann, H. W. and Machin, K. E. (1958). The mechanism of object location in Gymnarchus niloticus and similar fish. J. exp. Biol. 35, 451–486.

Lundberg, J. G., Lewis, W. M., JR., Saunders, James, F. [III] and Mago-Leccia, F. (1987). A Major Food Web Component in the Orinoco River Channel: Evidence from Planktivorous Electric Fishes. Science. 237, 81–83.

Markham, M. R., Ban, Y., McCauley, A. G. and Maltby, R. (2016). Energetics of Sensing and Communication in Electric Fish: A Blessing and a Curse in the Anthropocene? Integr. Comp. Biol. 56, 889–900.

Markham, M. R., McAnelly, M. L., Stoddard, P. K. and Zakon, H. H. (2009). Circadian and social cues regulate ion channel trafficking. PLoS biology. 7, e1000203.

Moulton, T. L. (2018). rMR: Importing Data from Loligo Systems Software, Calculating Metabolic Rates and Critical Tensions. https://CRAN.R-project.org/package=rMR.

Moulton, T. L., Chapman, L. J. and Krahe, R. (2020). Effects of hypoxia on aerobic metabolism and active electrosensory acquisition in the African weakly electric fish Marcusenius victoriae. J. Fish Biol. 96, 496–505.

Nakagawa, S., Schielzeth, H. and O’Hara, R. B. (2013). A general and simple method for obtaining R2 from generalized linear mixed-effects models. Methods Ecol. Evol. 4, 133–142.

Nelson, M. E. and MacIver, M. A. (1999). Prey capture in the weakly electric fish Apteronotus albifrons: Sensory acquisition strategies and electrosensory consequences. J. exp. Biol. 202, 1195–1203.

Pihl, L., Baden, S. P. and Diaz, R. J. (1991). Effects of periodic hypoxia on distribution of demersal fish and crustaceans. Mar. Biol. 108, 349–360.

Pörtner, H. O. and Peck, M. A. (2010). Climate change effects on fishes and fisheries: towards a cause-and-effect understanding. J. Fish Biol. 77, 1745–1779.

Pörtner, H. O. (2001). Climate change and temperature-dependent biogeography: Oxygen limitation of thermal tolerance in animals. Naturwissenschaften. 88, 137–146.

Pörtner, H. O. and Knust, R. (2007). Climate Change Affects Marine Fishes Through the Oxygen Limitation of Thermal Tolerance. Science. 315, 95–97.

Poulsen, S. B., Jensen, L. F., Nielsen, K. S., Malte, H., Aarestrup, K. and Svendsen, J. C. (2011). Behaviour of rainbow trout Oncorhynchus mykiss presented with a choice of normoxia and stepwise progressive hypoxia. J. Fish Biol. 79, 969–979.

Rahel, F. J. and Nutzman, J. W. (1994). Foraging in a lethal environment: Fish predation in hypoxic waters of a stratified lake. Ecology. 75, 1246–1253.

Reardon, E. E., Parisi, A., Krahe, R. and Chapman, L. J. (2011). Energetic constraints on electric signalling in wave-type weakly electric fishes. J. exp. Biol. 214, 4141–4150.

Salazar, V. L., Krahe, R. and Lewis, J. E. (2013). The energetics of electric organ discharge generation in gymnotiform weakly electric fish. J. exp. Biol. 216, 2459–2468.

Schmidtko, S., Stramma, L. and Visbeck, M. (2017). Decline in global oceanic oxygen content during the past five decades. Nature. 542, 335–339.

Serrano-Fernández, P. (2003). Gradual frequency rises in interacting black ghost knifefish, Apteronotus albifrons. J. Comp. Physiol. A Neuroethol. Sens. Neural. Behav. Physiol. 189, 685–692.

Stoddard, P. K. and Salazar, V. L. (2010). Energetic cost of communication. J. exp. Biol. 214, 200–205.

Wannamaker, C. M. and Rice, J. A. (2000). Effects of hypoxia on movements and behavior of selected estuarine organisms from the southeastern United States. J. Exp. Mar. Biol. Ecol. 249, 145–163.

Wolf, N. G. and Kramer, D. L. (1987). Use of cover and the need to breathe: The effects of hypoxia on vulnerability of dwarf gouramis to predatory snakeheads. Oecologia. 73, 127–132.

Yeager, D. P. and Ultsch, G. R. (1989). Physiological Regulation and Conformation: A BASIC Program for the Determination of Critical Points. Physiol. Zool. 62, 888–907.

